# Genetic characterization of chytrids isolated from larval amphibians collected in central and east Texas

**DOI:** 10.1101/451385

**Authors:** Thomas L. Marshall, Carlos R. Baca, Decio T. Correa, Michael R. J. Forstner, Dittmar Hahn, David Rodriguez

**Affiliations:** Department of Biology, Texas State University, 601 University Dr., San Marcos, TX 78666, USA; Department of Integrative Biology, University of Texas at Austin, 2415 Speedway, Austin, TX 78712, USA

**Author notes:** Corresponding author: Department of Biology, Texas State University, San Marcos, Texas, 78666, United States of America.

**Keywords:** *Batrachochytrium dendrobatidis*, pathogen dynamics, MLST, culture, genotyping, anuran host

## Abstract

Chytridiomycosis, an emerging infectious disease caused by the fungal pathogen *Batrachochytrium dendrobatidis* (*Bd*), has caused amphibian population declines worldwide. *Bd* was first described in the 1990s and there are still geographic gaps in the genetic analysis of this globally distributed pathogen. Relatively few genetic studies have focused on regions where *Bd* exhibits low virulence, potentially creating a bias in our current knowledge of the pathogen’s genetic diversity. Disease-associated declines have not been recorded in Texas (USA), yet *Bd* has been detected on amphibians in the state. These strains have not been isolated and characterized genetically; therefore, we isolated, cultured, and genotyped *Bd* from central Texas and compared isolates to a panel of previously genotyped strains distributed across the Western Hemisphere. We also isolated other chytrids not known to infect amphibians from east Texas. To identify larval amphibian hosts, we sequenced part of the COI gene. Among 37 *Bd* isolates from Texas, we detected 19 unique multi-locus genotypes, but found no genetic structure associated with host species, Texas localities, or across North America. Isolates from central Texas exhibit high diversity and genetically cluster with *Bd*-GPL isolates from the western U.S. that have caused amphibian population declines. This study genetically characterizes isolates of *Bd* from the south central U.S. and adds to the global knowledge of *Bd* genotypes.

## Introduction

Emerging infectious diseases (EIDs) — those that have recently increased in incidence, impact, virulence, geographic or host range, or have recently evolved — are a growing threat to human health and global biodiversity (Daszak *et al.* 2000, 2003). Over the past two decades, an increasing proportion of EIDs have been attributed to pathogenic fungi. These include powdery mildew on important crop plants, white-nosed syndrome in bats, snake fungal disease, colony collapse disorder in bees, and sea-fan aspergillosis in corals (Fisher *et al.* 2016; Fisher *et al.* 2012). Despite their immense economic and ecological impact, the genetic diversity of many of these fungi is still poorly understood (Fisher *et al.* 2016). None of these pathogens have been as devastating in their impact as *Batrachochytrium dendrobatidis* (*Bd*), the aquatic chytrid that induces chytridiomycosis, an epidermal infection in amphibians (Lips 2016).

*Bd*, a member of the Rhizophydiales order of fungi, causes infections when its flagellated zoospores encyst in a host’s skin and develop into zoosporangia (Letcher *et al.* 2006; Longcore *et al.* 1999). It was first described and attributed to disease outbreaks in the late 1990s (Longcore *et al.* 1999) and is now believed to be responsible for declines in amphibian populations dating back at least to the 1970s (Lips *et al.* 2004). *Bd* is found on every continent except Antarctica, is known to have infected over 700 species of amphibians, and has caused mass mortality events including species extinctions in Central and northern South America, the western U.S., Australia, and Spain (Berger *et al.* 1998; Bosch *et al.* 2001; Lips *et al.* 2006; Vredenburg *et al.* 2010). However, certain amphibian species and geographic regions have seemingly been unaffected by chytridiomycosis, even where *Bd* is present (Lips 2016).

Recent phylogenomic analyses of *Bd* have revealed multiple lineages that include the widespread global panzootic lineage (GPL), which is implicated in most disease outbreaks and can be further divided into a temperate clade (*Bd*-GPL1) and a tropical clade (*Bd*-GPL2). Other lineages include the putatively endemic and less virulent *Bd*-Cape (found in South Africa and Spain), *Bd*-CH from Switzerland (Farrer *et al.* 2011), and *Bd*-Korea (Bataille *et al.* 2013). Sampling efforts have also led to the discovery of a strain in Japan that may be host specific (Goka *et al.* 2009) and *Bd*-Brazil (Jenkinson *et al.* 2016; Rodriguez *et al.* 2014; Schloegel *et al.* 2012), which genomic analyses have indicated is the earliest diverged *Bd* lineage currently known (Rosenblum *et al.* 2013).

Recently revealed diversity within the *Bd* clade, coupled with the fact that there are still several geographic gaps in our current knowledge of *Bd* genotypes (James *et al.* 2015), suggests additional diversity is yet to be discovered. However, relatively few studies have examined *Bd* genotypes from regions lacking population declines, potentially creating a bias in our knowledge of the pathogen’s genetic diversity (Bataille *et al.* 2013). It is likely that *Bd*’s origin can be traced to a region in which amphibians have co-evolved with the fungus and where severe population declines and extinctions are not occurring (James *et al.* 2015). While recent work has begun to explore such regions of potential origin, genotyping in additional understudied areas is imperative and is facilitated by isolation and culturing of the fungus (Bataille *et al.* 2013; Jenkinson *et al.* 2016).

In the U.S., *Bd*-GPL has shown patterns of introduction and spread (Padgett-Flohr & Hopkins 2009) and has caused host declines in the Sierra Nevada of California (Briggs *et al.* 2010; Wake & Vredenburg 2008). On the other hand, long-term persistence and a lack of chytridiomycosis related host declines is also evident in the Midwest (Illinois) (Talley *et al.* 2015). Texas is a part of the south-central U.S. where, like the Midwest, no known chytridiomycosis-associated amphibian declines or extinctions have been detected. Studies have confirmed the presence of *Bd* in the eastern (Saenz *et al.* 2010) and central (Gaertner *et al.* 2012; Gaertner *et al.* 2009a; Gaertner *et al.* 2009b; Gaertner *et al.* 2010) parts of the state, yet no strains from these regions have been genotyped or isolated thus far (James *et al.* 2015). While a lack of known *Bd*-related population declines or extinctions in the area could be an indication that the pathogen has a long local history, other factors, such as climate or host resistance, may be responsible for attenuating disease outbreaks in the region. Studying the genetics of the fungus in Texas and its amphibian hosts is an important first step in understanding potential disease dynamics between amphibians and *Bd* in the region. Additionally, by genetically characterizing isolates from Texas and placing them in the context of a more broadly distributed panel of strains, we can begin to gain a more complete evolutionary picture of the *Bd* group, and perhaps a better understanding of the origins of its most virulent lineages. Therefore, the goal of this study was to isolate *Bd* from larval anurans in central and east Texas, genetically identify anuran hosts, and compare genetic variation among chytrid strains present in this region to those distributed across the Western Hemisphere.

## Methods and Materials

### Host collection

We collected anuran larvae from ponds in Travis, Hays, Bastrop, Kaufman, and Houston counties in central and eastern Texas from December 2015 to April 2016 (collection permit: SPR-0102-191) and stored them in 50 ml Falcon tubes or new plastic containers with water from the collection site (Table 1). We transported all collected tadpoles, in Styrofoam containers kept cool with synthetic ice packs, to Texas State University where they were euthanized following animal care and use protocols IACUC2014116499 and IACUC2015121220. Each tadpole was given a unique identifier, and a tail clipping from each tadpole was preserved in 95% ethanol for DNA extraction.

**Table 1.** Collection sites (abbreviations) for anuran larvae, collection dates, county (Texas, USA), GPS coordinates of specific localities, number of anuran larvae collected, number of chytrid strains isolated (including non-*Batrachochytrium dendrobatidis* chytrids), and host species from which chytrids were successfully isolated.

| Site | Date | County | Latitude (N) | Longitude (W) | Larvae Collected | Chytrid Isolates | Host Species |
| --- | --- | --- | --- | --- | --- | --- | --- |
| Abby's Pond (AP) | Feb 2016 | Hays | 29.9772 | -97.8984 | 6 | 3 | *R. berlandieri* |
| Bee Cave (BC) | Mar 2016 | Travis | 30.2542 | -97.9392 | 6 | 4 | *R. berlandieri* |
| Brackenridge Field Laboratory (BFL) | Mar 2016 | Travis | 30.2837 | -97.7801 | 40 | 11<sup>a</sup> | *R. berlandieri* |
| Davy Crockett National Forest (DCNF) | Feb 2016 | Houston | 31.4083 | -95.1667 | 21 | 3<sup>b</sup> | *R. clamitans* |
| Griffith League Ranch (GLR) | Dec 2015 | Bastrop | 30.2148 | -97.2576 | 6 | 0 | *R. sphenocephala* |
| Lakeway (LW) | Mar 2016 | Travis | 30.3766 | -98.0421 | 6 | 6 | *R. berlandieri* |
| RRR Ranch (RRR) | Mar 2016 | Kaufman | 32.5332 | -96.4880 | 47 | 13 | *R. sphenocephala* |
| Wimberley (WM) | Mar 2016 | Hays | 29.9955 | -98.2163 | 8 | 1 | *R. berlandieri* |
| **Total** |  |  |  |  | 140 | 41 |  |

<sup>a</sup> includes 1 non-*Bd* isolate
<sup>b</sup> all non-*Bd* isolates

### Fungal isolation, culture, and cryopreservation

We excised the keratinized jaw sheaths and tooth rows from each tadpole and transferred them to 1% typtone agar petri dishes. We cleaned the mouthparts by dragging them through fresh agar and transferring them to 1% tryptone agar petri dishes with 200 mg/ml penicillin-G and 200 mg/ml streptomycin sulfate for isolation of chytrids (Longcore 2000). We monitored the plates for growth of zoosporangia and active zoospores for up to two weeks. Once enough active zoospores were observed, we removed a section of the agar containing active zoospores to tryptone broth (16 g tryptone in 1,000 ml deionized water; autoclaved) to cultivate the isolate. Once there was a high density of active zoospores, we transferred a 1-ml aliquot of broth from each isolate to a 1.5-ml centrifuge tube. We used a cryoprotectant solution consisting of 80 ml tryptone broth, 10 ml dimethyl sulfoxide, and 10 ml fetal calf serum to preserve all chytrid isolates (Boyle *et al.* 2003). For each isolate, we briefly centrifuged (2,000 rpm) tryptone broth cultures in 1.5 ml cryotubes, discarded the supernatant, and added 600 µl of cryoprotectant. All isolates were given a serially numbered unique identifier (TXST – Texas State University) and were stored at −80°C (Appendix I).

### Genetic methods

To extract DNA, we centrifuged 1 ml aliquots of live broth cultures, discarded the supernatant, and extracted DNA from the pellet using the Mammalian Tissue and Rodent Tail Genomic DNA Purification Protocol in the Thermo Scientific GeneJET Genomic DNA Purification Kit #K0722 (Thermo Fisher Scientific, Inc.). We determined extraction success with gel electrophoresis using a 1% agarose gel stained with GelRed (Biotium) in 1X TBE buffer. We performed conventional Polymerase Chain Reaction (PCR) on DNA from each successfully extracted isolate using primers for previously described multilocus sequence typing (MLST) markers (8009X2, BdC5, BdSc2.0, BdSc4.16, BdSc7.6, R6046, BdSC6.15) and the 18S rRNA gene (James *et al.* 2009; Jenkinson *et al.* 2016; Morehouse *et al.* 2003; Morgan *et al.* 2007; Schloegel *et al.* 2012) (Table S1). Amplifications were performed in 25 µl volumes consisting of 12.5 µl DreamTaq PCR Master Mix (2X) (Thermo Fisher Scientific, Inc.), 11.5 µl nuclease free water, 0.25 µl forward primer (10 µM), 0.25 µl (10 µM) reverse primer, and 0.5 µl template DNA. Thermocycling conditions consisted of an initial denaturing step of 2 min at 95°C, then 32 cycles of 30 s at 95°C, 30 s at 52°C to 60°C depending on the primer pair (Table S1), 45 s at 72°C, with a final extension of 10 min at 72°C. We treated 5 µl aliquots of PCR product with 2 µl ExoSAP-IT (Affymetrix Inc.) and incubated the total volume at 37°C for 15 min and then 80°C for 15 min. Using each of the MLST primers, we performed cycle sequencing reactions using Big Dye v3.1 dye terminator (Applied Biosystems, Inc.). We incubated hydrated G-50 Sephadex (2.6 g/45 ml H_2_O) at room temperature for 30 min, pipetted 400-µl aliquots into individual wells in a filter plate, and centrifuged the plate at 3000 rpm for 2 minutes to create a matrix through which cycle sequenced products were passed for purification. We dehydrated the purified cycle sequenced products, added 12 µl of formamide, and incubated them for 3 min at 94°C and immediately cooled them to 4°C. We electrophoresed the cycle sequenced products on an ABI 3100-Avant genetic analyzer (Applied Biosystems, Inc.) and trimmed and edited the resulting chromatograms using Geneious v9.1.5 (Kearse *et al.* 2012).

### Data Analysis

We genotyped strains using reference sequences from Jenkinson *et al.* (2016). We then used a set of genotyped strains from across North and South America to perform comparative analyses and test for genetic structure across the Western Hemisphere. We limited our analyses to the Western Hemisphere owing to the relatively few number of strains sequenced at these same markers in the Eastern Hemisphere. These strains were selected from two previous studies that used at least six of the same genetic markers used in our study (Jenkinson *et al.* 2016; Schloegel *et al.* 2012). We excluded any strains that had been isolated from amphibians not found in the wild (this included animals from zoos, farms, and markets) for our analyses. The resulting dataset consisted of 177 strains, including the 37 Texas strains isolated in our study. Before calculating measures of genetic diversity, we divided our dataset into populations and subpopulations. These populations (and subpopulations) included temperate America (subpopulations: eastern North America, Texas, and western North America), tropical America (subpopulations: Panama, Brazil-GPL), *Bd*-Brazil, and *Bd*-Brazil/GPL hybrids. For each population and subpopulation, we calculated genotypic diversity and allelic richness using the R package POPGENREPORT (Adamack & Gruber 2014). Next, we clone-corrected our datasets by removing identical genotypes from the same region. This resulted in a dataset that included the genotypes of 68 isolates, including 15 unique Texas isolates, at six MLST markers (Appendix 2). We then calculated gene diversity, or expected heterozygosity (*H*_E_), and observed heterozygosity (*H*_O_) using the R package ADEGENET (Jombart 2008) and calculated pairwise F_ST_ values using GENEPOP (Rousset 2008).

To test for genetic structure among *Bd* isolates, we used STRUCTURE (Pritchard *et al.* 2000) with values of *K* (number of genetic demes defined *a priori*) ranging from 1 to 12 and 5 iterations per *K*, using 500,000 Markov Chain Monte Carlo repetitions with a burn-in of 100,000. We determined the best number of populations by calculating ⋄*K* (Evanno *et al.* 2005) using STRUCTURE HARVESTER (Earl & Vonholdt 2012) and obtained average membership probabilities across all iterations per *K* using the program CLUMPP (Jakobsson & Rosenberg 2007). We performed an initial analysis with only our Texas isolates and their genotypes at all seven loci (8009×2, BdC5, BdSC4.16, BdSC7.6, R6046, BdSC6.15, BdSC2.0) to test for genetic structure within the state, followed by a second analysis combining the Texas isolates with the set of isolates from North and South America that had been sequenced and genotyped using at least six of the same markers (8009×2, BdC5, BdSC4.16, BdSC7.6, R6046, BdSC6.15). Both STRUCTURE analyses were performed using clone-corrected data.

### Host Sequencing and Identification

We extracted DNA from tail muscle of collected tadpoles and toes of seven adult *Rana berlandieri* using the same extraction protocol described for *Bd*. Conventional PCR was performed to amplify the cytochrome c oxidase I (COX1) gene (Table S1), and PCR products were purified and sequenced using the same methods described for *Bd* sequencing. Chromatograms were trimmed, edited and aligned in Geneious v 9.1.5 (Kearse *et al.* 2012). Using Clustal W, implemented through Geneious, we carried out a multiple sequence alignment of host sequence data (cost matrix = clustalw, gap open cost = 15, gap extend cost = 6.66). We used sequence similarity to assign anuran larvae to species and visualized the distance matrix (HKY) using a neighbor-joining topology. We included reference sequences from *Pseudacris streckeri* (GenBank accession no. **KJ536156**), *Rana muscosa* (GenBank accession no. **KU985709**), *R. clamitans* (GenBank accession no. **KY587195**), and *R. sphenocephala* (GenBank accession no. **KT388406**) and newly generated reference sequences for adult *R. berlandieri* (GenBank accession no. **MG969220** - **MG969226**). We suspected that some of the larvae belonged to these species based on geographic distribution and general morphology. *Rana muscosa* was chosen as an outgroup to Texas ranids.

## Results

We attempted to isolate *Bd* from a total of 140 tadpoles collected from sites in central, east, and north-central Texas (Fig. 1). From this total, we successfully cultured 41 chytrid isolates from 40 individual tadpoles comprising three host species. Of these 41 isolates, 37 were identified as *Bd* (Appendix 1). A total of 28 isolates have been characterized using all seven MLST markers, seven have been characterized at six loci, and two were characterized at five loci (Supplemental data). We were unable to amplify DNA of four isolates from two *R. clamitans* larvae using these markers but did amplify and sequence part of the 18S rRNA gene (Appendix 1). Using the NCBI BLAST search tool (Clark *et al.* 2016), we matched isolates TXST038 and TXST039 to *Rhizophlyctis harderi* (GenBank accession no. **AF164272**; match = 98%, coverage = 100%, E value = 0.0), which has recently been re-named *Uebelmesseromyces harderi* (Powell *et al.* 2015), TXST041 to *Chytriomyces* sp. (GenBank accession no. **DQ536486**; match = 98%, coverage = 100%, E value = 0.0), and isolate TXST038 to *Hyaloraphidium curvatum* (GenBank accession no. **NG_017172**; match = 99%, coverage = 100%, E value = 0.0), all of which are species of chytrids not known to infect vertebrate hosts. Among the 37 *Bd* isolates, there were 19 unique multi-locus genotypes (MLGs), and no MLGs were found at more than one collection site (Supplemental data).

**Fig. 1.**
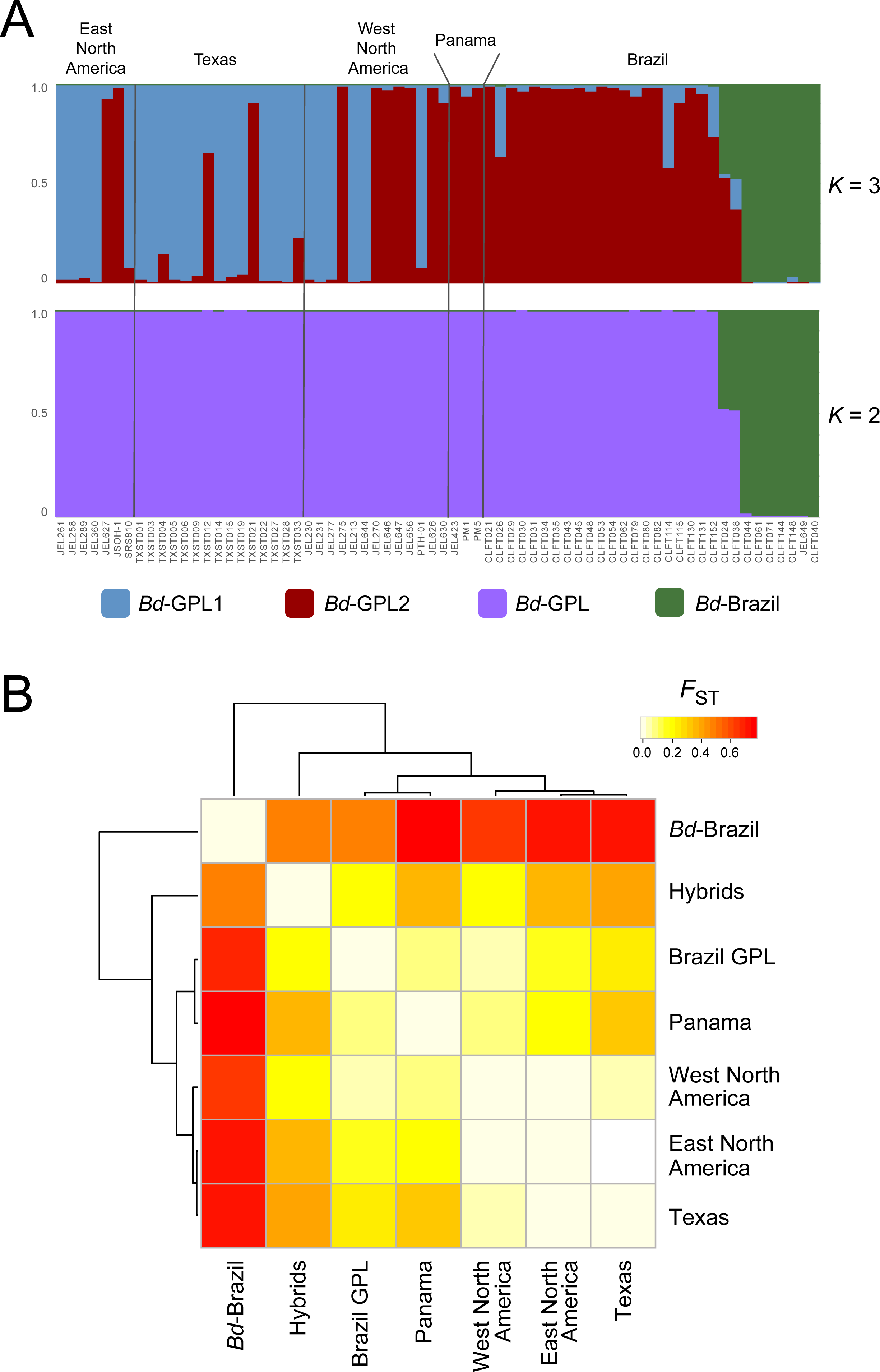
Sites where anuran larvae were collected and the proportion of chytrid isolates genetically identified to a known *Bd* strain or other chytrid. The background colors indicate mean maximum temperature in the region (USDA/NRCS - National Geospatial Center of Excellence 2015) with the optimal temperature range for *Bd* growth demarcated in the legend (Piotrowski *et al.* 2004).

A neighbor-joining topology (Fig. S1) revealed four distinct clades among potential hosts. Three clades were associated with *R. clamitans, R. sphenocephala,* and *Pseudacris streckeri* reference sequences. We suspected that some of our tadpole specimens were *Rana berlandieri*, although a COI reference sequence for this species was unavailable in GenBank. Thus, we sequenced seven *R. berlandieri* metamorphs from central, south, western parts of Texas (Appendix 1) to verify species assignments. All newly generated sequences were accessioned into GenBank (chytrid 18S, **MG979804–MG979842**; amphibian COI, **MG969220– MG969347**).

### Measures of Genetic Diversity

Gene diversity (*H*_E_) among the clone corrected Western Hemisphere dataset was 0.506 and observed heterozygosity (*H*_O_) was 0.370. When all *Bd*-GPL strains were divided into temperate and tropical groups, the temperate group had both higher *H*_E_ and *H*_O_ (0.402 and 0.394, respectively) than the tropical group (0.358 and 0.340, respectively). Both groups had higher *H*_E_ and *H*_O_ than *Bd*-Brazil (0.189 and 0.214, respectively). While larger pooled populations had higher expected than observed heterozygosity, the subpopulations in our dataset tended to show the reverse trend (except for the GPL in western North America and southeastern Brazil). While Texas strains had lower *H*_E_ (0.348) than both eastern and western North American subpopulations, *H*_O_ (0.387) was intermediate between the two other North American regions. All North American subpopulations, including Texas, had higher *H*_O_ than Panama and southeastern Brazil, while Texas was the only North American subpopulation to have lower *H*_E_ than the GPL in southeastern Brazil. Panama had the lowest *H*_E_ and *H* _O_of all GPL subpopulations. The temperate GPL had higher genotypic diversity (0.491) and allelic richness (1.81) compared to the tropical GPL (0.250 and 1.68, respectively) and *Bd*-Brazil (0.280 and 1.34, respectively). Texas strains had lower genotypic diversity (0.405) and allelic richness (1.72) than eastern and western North American strains but displayed greater diversity in these measures than both tropical groups. Not surprisingly, the GPL/*Bd*-Brazil hybrid strains had higher measures of genetic diversity than all other groups (Table 2).

**Table 2.** Genetic diversity of *Batrachochytrium dendrobatidis* isolates collected in Texas compared to other isolates distributed across the Western Hemisphere. Genetic diversity indices were calculated by averaging across the same six MLST markers.

| Region/Population | N<sup>a</sup> | MLG<sup>b</sup> | Genotypic Diversity | Mean Allelic Richness | Clone Corrected Data |  |
| --- | --- | --- | --- | --- | --- | --- |
|  |  |  |  |  | Observed Heterozygosity | Expected Heterozygosity |
| Temperate America (*Bd*-GPL) | 57 | 28 | 0.491 | 1.81 | 0.394 | 0.402 |
| Eastern N.A. | 7 | 7 | 1.000 | 1.83 | 0.429 | 0.378 |
| Western N.A. | 13 | 13 | 1.000 | 1.91 | 0.385 | 0.435 |
| *Texas* | 37 | 15 | 0.405 | 1.72 | 0.387 | 0.348 |
| Tropical America (*Bd*-GPL) | 92 | 23 | 0.250 | 1.68 | 0.34 | 0.358 |
| Panama | 3 | 3 | 1.000 | 1.59 | 0.333 | 0.259 |
| Southeastern Brazil | 89 | 21 | 0.236 | 1.67 | 0.341 | 0.354 |
| **All *Bd*-GPL** | **149** | **47** | **0.315** |  | **0.373** | **0.419** |
| Hybrids (GPL and *Bd*-Brazil) | 3 | 2 | 0.667 | 1.96 | 0.833 | 0.438 |
| ***Bd*-Brazil** | **25** | **7** | **0.280** | **1.34** | **0.214** | **0.189** |
| **Western Hemisphere** | **177** | **56** | **0.316** |  | **0.370** | **0.506** |

<sup>a</sup> Sample size. <sup>b</sup> Number of multi-locus genotypes

### Cluster Analyses

We performed a STRUCTURE analysis using seven loci on the 19 unique MLGs from Texas and did not detect population structure among these strains. We then conducted a STRUCTURE analysis on the combined dataset that included our Texas isolates, as well as additional Western Hemisphere genotypes from two previous studies (Jenkinson *et al.* 2016; Schloegel *et al.* 2012). For these analyses, we chose six MLST markers (r6046, BdSc6.15, 8009x2, BdC5, BdSC4.16, BdSC7.6) that had been sequenced in all three studies, thus we used 15 unique MLGs from Texas to compare against other New World isolates (Appendix 2).

Our STRUCTURE analysis indicated two to three genetically distinct demes in our dataset, with the highest ⋄ *K* value occurring at *K*= 2 and second highest value at *K*= 3 (Fig. S2). By using the already genetically identified isolates from the two previous studies as references, we determined that at *K*= 2, our clusters corresponded to *Bd*-GPL and *Bd*-Brazil, while at *K*= 3 substructure within the GPL corresponded to the two major clades within this group; namely *Bd-* GPL1 and *Bd*-GPL2 (Fig. 2A). However, we did not detect the presence of any *Bd*-Brazil strains in Texas. We set an arbitrary threshold of *q* = 0.80 for assigning strains to a particular cluster. Ambiguous strains that did not show strong support for a cluster included the putative GPL/*Bd-* Brazil hybrids, and two strains from Texas and three from Brazil that exhibited mixed support for the two GPL lineages. Pairwise *F*_ST_ values indicate much greater differentiation between *Bd-* Brazil and all other populations, while Texas isolates show no difference from eastern North American isolates and relatively low differentiation (0.040) from isolates in western North America (Fig. 2B, Table S2).

**Fig. 2.**
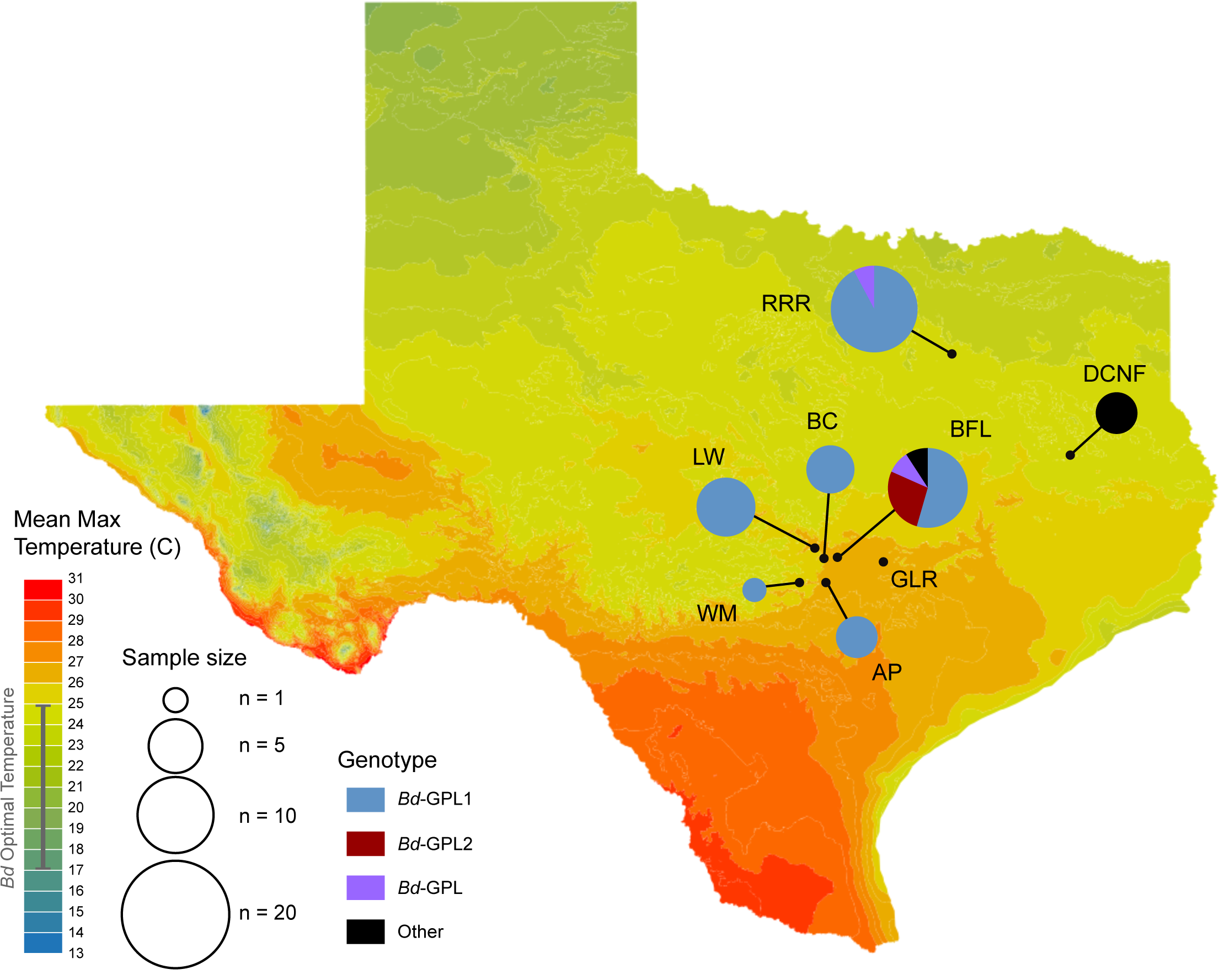
(**A**) STRUCTURE results for *K =*2 and *K =*3 averaged from five iterations using CLUMPP. Assignment probabilities to the clusters are represented on the *y-* axis, and each bar represents one of the 68 isolates from a six MLST dataset, including 15 isolates collected for this study and 53 additional strains from Schloegel et al. (2012) and Jenkinson et al. (2016). (**B)**Heat map of pairwise *F*_ST_ values by origin calculated from the dataset of 68 *Bd* isolates, including 15 isolates from central Texas and 53 additional strains from Schloegel et al. (2012) and Jenkinson et al. (2016).

## Discussion

*Bd* has shown contrasting pathogen dynamics in North America (Briggs *et al.* 2010; Padgett-Flohr & Hopkins 2009; Talley *et al.* 2015; Wake & Vredenburg 2008), yet genetic information from cultured isolates is mainly restricted to the western and eastern U.S., leaving a large swath of the central U.S. without data on *Bd* genotypes, and thus information on strain types. We successfully isolated 37 chytrids identified as *Bd* and four other chytrids (three species) from three different amphibian host species native to central and east Texas. As a population, *Bd* isolates in this region do not show significant genetic differentiation from other North American isolates. Further, all *Bd* isolates collected in this study belong to the globally distributed *Bd*-GPL, and none show genetic affinities to other major lineages (e.g. *Bd*-Brazil, *Bd-* CH, *Bd*-CAPE, or *Bd*-Korea).

Our measures of genetic diversity across geographic regions in the Western Hemisphere are congruent with previous studies. James *et al.* (2009) found lower *H*_E_ and *H*_O_ among tropical GPL strains compared to temperate GPL strains using MLST markers. The latitudinal gradient in genetic diversity observed by Velo-Anton *et al.* (2012) from California to Panama is also congruent with our estimates. While North American strains have the highest measures of genetic diversity, Texas strains, which represent the southernmost group of North American strains in our dataset, tend to have slightly lower measures of diversity compared to the rest of the continent. Meanwhile, strains from Panama, the region closest to the equator in our dataset, have the lowest measures of diversity among the GPL. These trends could be reflective of a longer history of *Bd* in North America compared to the tropics, but we stress the need for further isolations from North America and genomic sequencing of these strains to draw stronger conclusions about the demographic history of *Bd* in the Western Hemisphere.

The STRUCTURE results show support for two genetic demes within the GPL in Texas and throughout North America. These clusters, which are congruent to *Bd*-GPL1 and *Bd*-GPL2 (Rosenblum *et al.* 2013), are not structured by geography or host species. Of the 37 cultured Texas isolates, 32 appear to cluster with *Bd*-GPL1 isolates. Three Texas isolates (TXST021, TXST026, and TXST035) cluster with *Bd*-GPL2, and two isolates (TXST012 and TXST033) exhibit some degree of assignment uncertainty between both *Bd*-GPL groups (Fig. 2A). Evidence for hybridization between *Bd* lineages has been detected (Jenkinson *et al.* 2016; Schloegel *et al.* 2012), thus it is possible that these ambiguous strains could be the result of hybridization between the two *Bd*-GPL lineages. The presence of the *Bd*-GPL2, a mainly tropical clade (Rosenblum *et al.* 2013), in Texas is not a surprising discovery given that *Bd*-GPL2 isolates have also been documented in isolated locations across North America, but its apparent rarity relative to the *Bd*-GPL1 is consistent with the pattern seen across the continent (James *et al.* 2015). We should point out, however, that there is some ambiguity in assigning membership to these two sub-lineages within the GPL.

At the continental scale, *Bd* isolates in Texas genetically cluster with isolates responsible for severe outbreaks of chytridiomycosis in the western United States (Fig. 2B). This finding leaves us with a still unanswered question—why do these strains, which are genetically similar, cause disease and mortalities in some regions but not others? If the pathogen is genetically similar in these different regions, the other two points of the epidemiological triangle—host and environment—should be explored further. One intuitive explanation for the lack of known chytridiomycosis outbreaks in Texas is the influence of climate. Piotrowski *et al.* (2004) reported *Bd*’s optimal temperature range to be 17-25°C with a maximum threshold of 28°C. In Texas, amphibians may be able to clear or reduce *Bd* infections before they become lethal coincident to hot summers, when temperatures regularly exceed 32°C (Nielsen-Gammon 2011). Indeed, *Bd* infections in *Acris crepitans* populations in central Texas have revealed a seasonal pattern in infection intensity, with peaks occurring in early spring followed by a decline throughout the summer months (Gaertner *et al.* 2012; Gaertner *et al.* 2009b). Additionally, temperatures in most parts of the state regularly exceed *Bd*’s thermal optimum for over six months of the year (U.S. climate data 2017). However, weather conditions cannot explain the lack of *Bd*-associated morbidity in central Texas salamanders of the genus *Eurycea*, which dwell in cool, thermally stable aquifers and springs. While *Bd* infections in these salamanders have been documented (Gaertner *et al.* 2009a), symptoms of chytridiomycosis have not been detected in these populations despite several years of investigation (Bendik 2017; Bowles *et al.* 2006; Pierce *et al.* 2010). Differences in host resistance among and even within species have been documented (Savage & Zamudio 2011; Woodhams *et al.* 2007), so it is possible that host species immunity plays a role in limiting the severity of disease outcomes in central Texas. The *Bd* strains isolated for this study will facilitate future studies necessary to disentangle the effects of both climate and host resistance on the dynamics of *Bd* infection in this region.

Our isolation and culturing efforts also recovered chytrids not known to infect vertebrate hosts, namely *Uebelmesseromyces harderi* (Rhizophydiales)*, Chytriomyces* sp. (Chytridiales), and *Hyaloraphidium curvatum* (Monoblepharidales) (Fig. 1, Appendix 1). Zoosporic fungi of the phylum Chytridiomycota are generally saprobes or parasites of algae, plants, and invertebrates (Barr 2001). It is possible that these chytrids were present in the pond water from which these tadpoles were collected, and that they somehow persisted on the tadpole mouthparts despite our following established procedures to clean them by dragging through sterile agar several times (Longcore *et al.* 1999). Although studies of the morphology and systematics of *U. harderi*, *Chytriomyces*, and *H. curvatum* can be found in the literature, little appears to be known about the life history of these species (Forget *et al.* 2002; Letcher & Powell 2002; Powell *et al.* 2015; Ustinova *et al.* 2000). Many chytrids have similar, uninformative morphological features (James *et al.* 2000), and what appear to be successful isolations may not represent *Batrachochytrium* species. Yet, these isolations and cultures are still useful in diversity discovery efforts, especially among the Chytridiomycota, a relatively understudied group of fungi (James *et al.* 2000). By sequencing isolates at the 18S rRNA gene, we were able provide genetic and locality reference data for these species. Owing to the lack of data for environmental chytrids, we recommend genetic characterization of isolated cultures using general primers before—rather than in response to—unsuccessful amplifications of *Bd* specific primers. These efforts will inform genetic databases and accurate curation of cryopreserved microscopic fungal collections.

Our investigation of genetic variation among *Bd* isolates from parts of Texas is an important first step in understanding disease dynamics between amphibians and *Bd* in this understudied region. We have gained a more complete picture of the distribution of *Bd* strains in this region and have begun to fill a large sampling gap for *Bd* isolates in North America. There are also important conservation implications to our understanding of chytridiomycosis dynamics in the region as well. Texas is home to 16 endemic amphibian species, many of which are federally listed as species of concern, and all of which are found in either the Edwards Plateau or the coastal prairies of Texas (Tipton *et al.* 2012). Knowledge of the *Bd* strains present in these regions can inform assessments of potential disease risks to host species in the face of continued climate change and habitat alteration.

## Acknowledgments

We thank Abigail Romo, Charles and Rebecca Raleigh, and the Brackenridge Field Laboratory for access to collection sites. We dedicate this work to the memory of Jonathan Raleigh. We thank Dan Saenz, Trina Guerra, Stephen Roussos, Jeremy Weaver, Brandon Cranup, William Keitt, Jacob Owen, Madison Torres, Andrea Villamizar, Stephen Harding, Carlos Baca, Daniel Puckett, Edith Perez, Devlin Jackson, Anna Gates, and Stephanie Toussaint for assistance in the lab and field. We thank Ana V. Longo for reviewing an earlier version of this manuscript. This work was funded by a National Science Foundation Graduate Research Fellowship to TLM (DGE-1144466), a Texas State University Research Enhancement Program grant to DR, Start-up funds from Texas State University to DR, and a CAPES fellowship (BEX 1176/13-7) and Texas EcoLab Program grant to DTC.

## APPENDICES

**Appendix 1.**
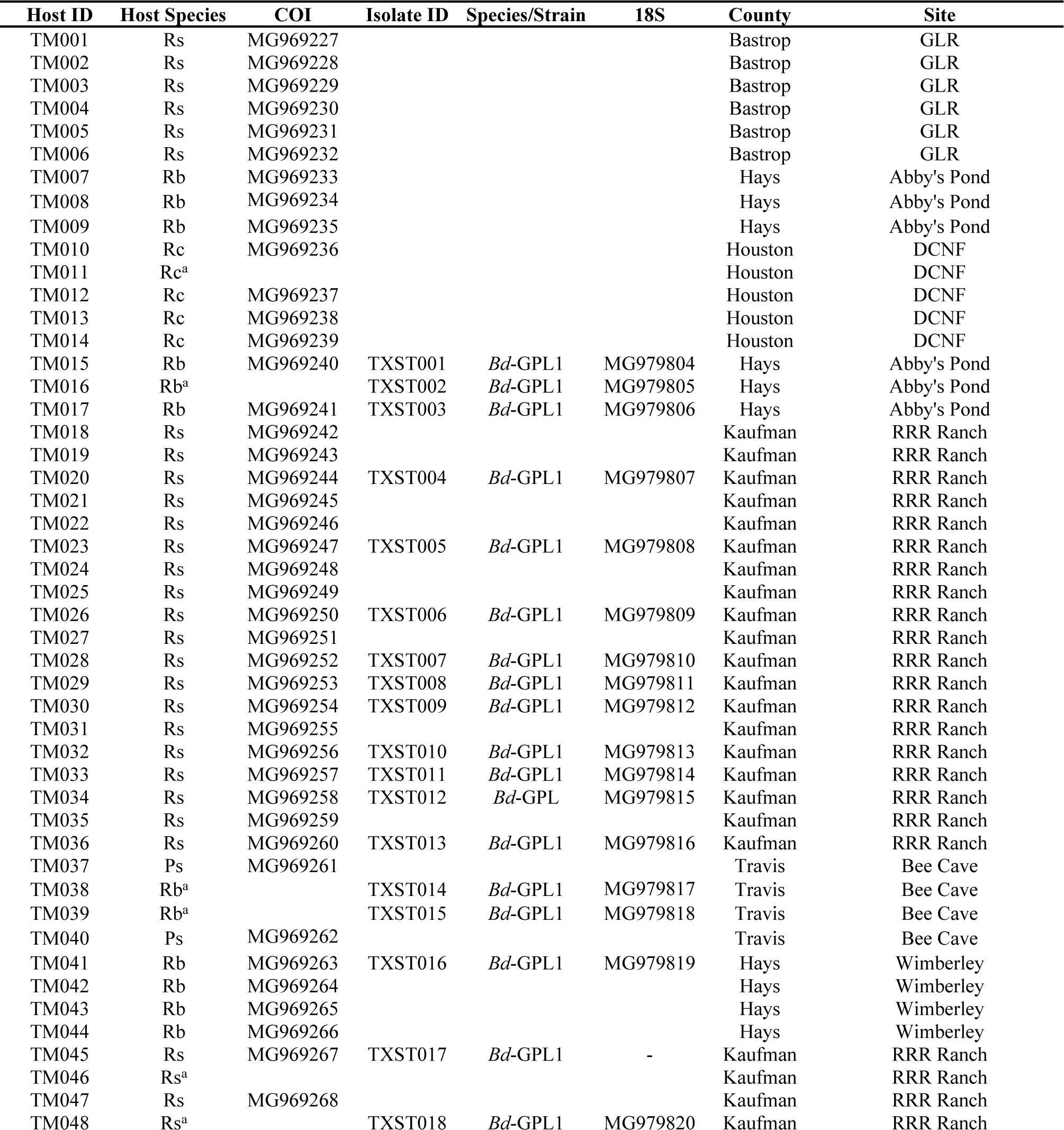
Amphibian larvae and adults sampled for this study with collector number (TM = Thomas Marshall, MF = Michael Forstner Tissue Catalog), host species (*Rs* = *Rana sphenocephala*, *Rb* = *R. berlandieri*, *Rc* = *R. clamitans, Ps = Pseudacris streckeri*), host COI GenBank accession number, isolate number, chytrid taxon (*Bd = Batrachochytrium dendrobatidis, Uh =Uebelmesseromyces harderi, Ch = Chytriomyces* sp.*, Hc = Hyaloraphidium curvatum*), chytrid 18S GenBank accession number, and locality information.

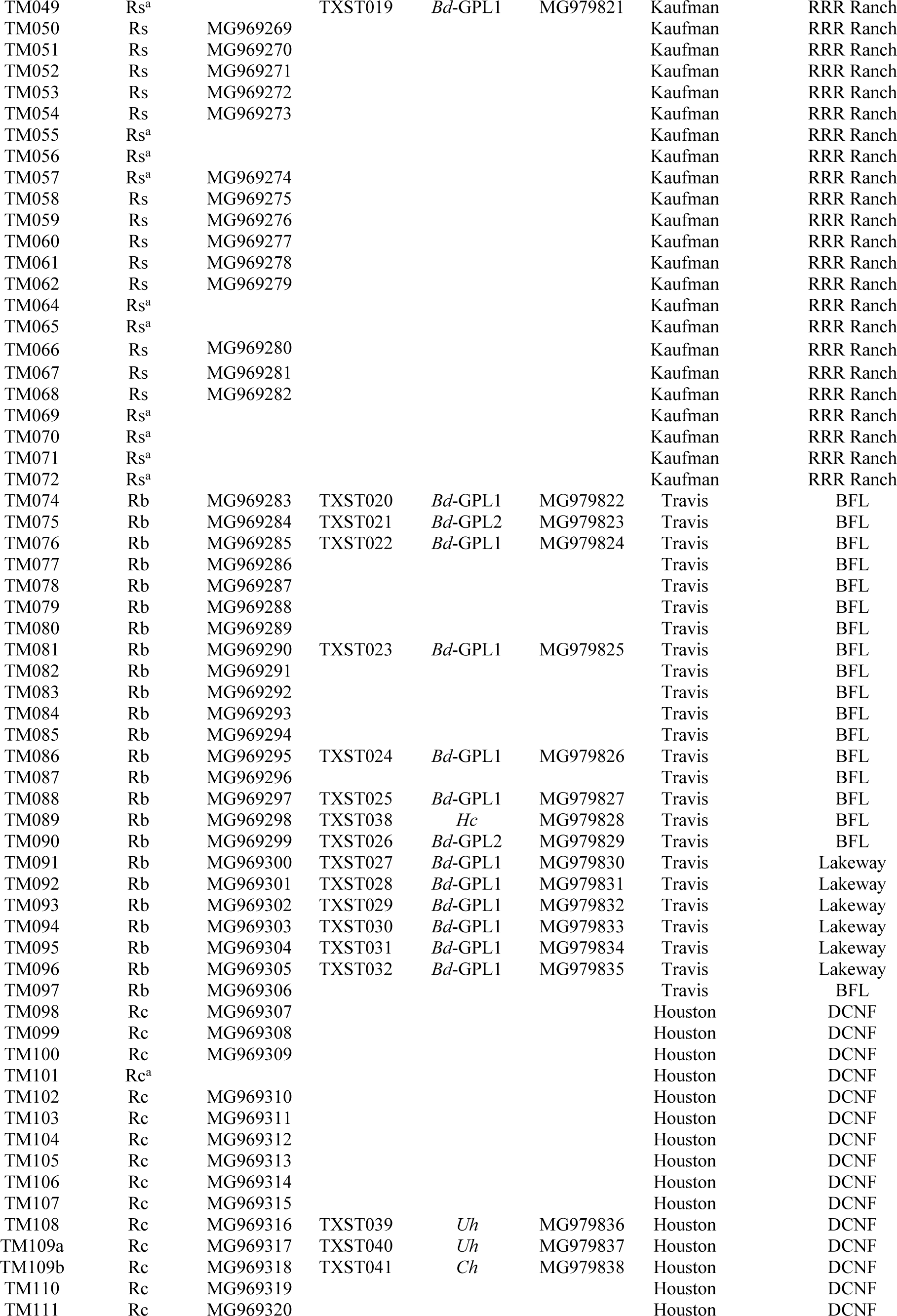

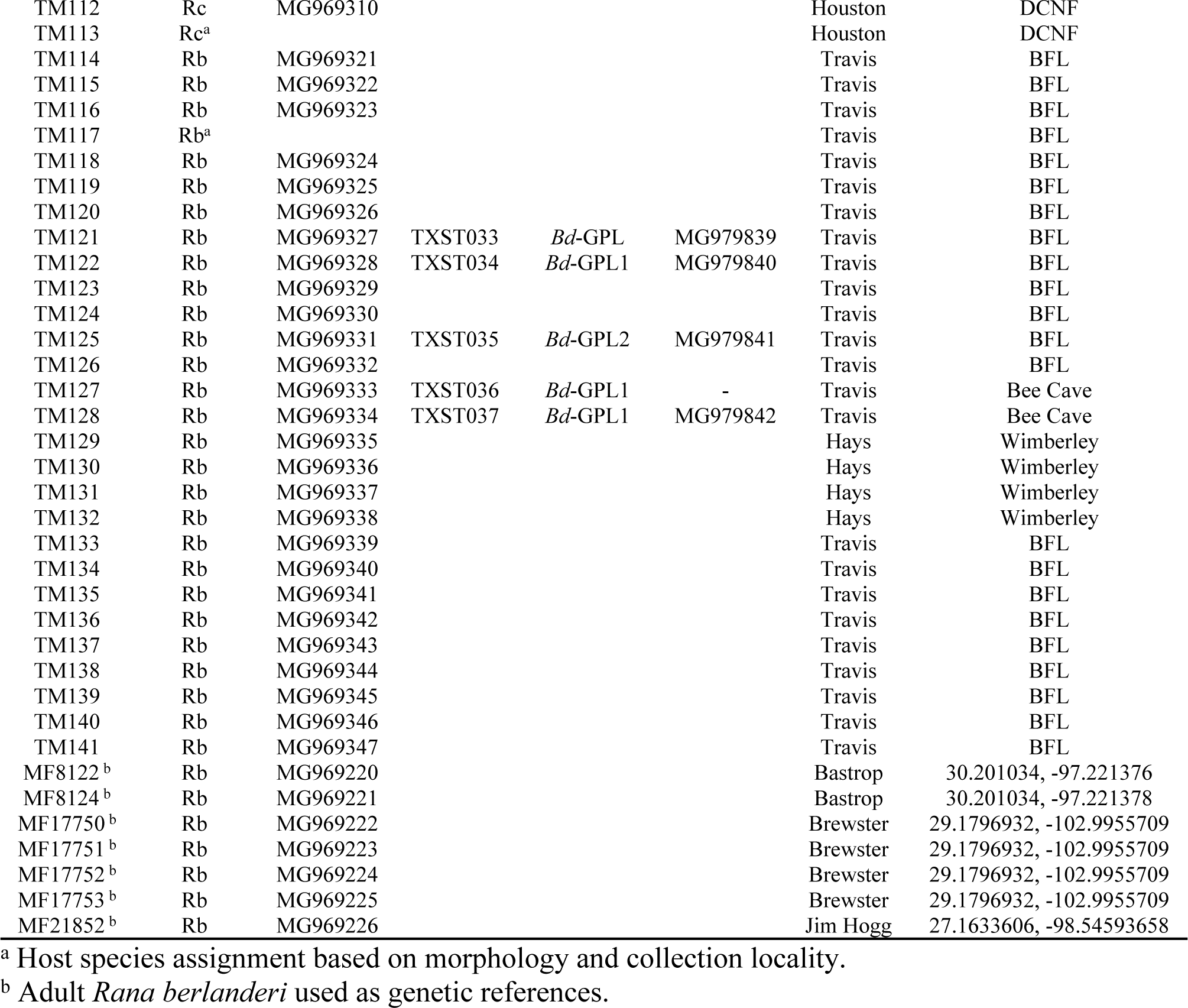

**Appendix 2.**
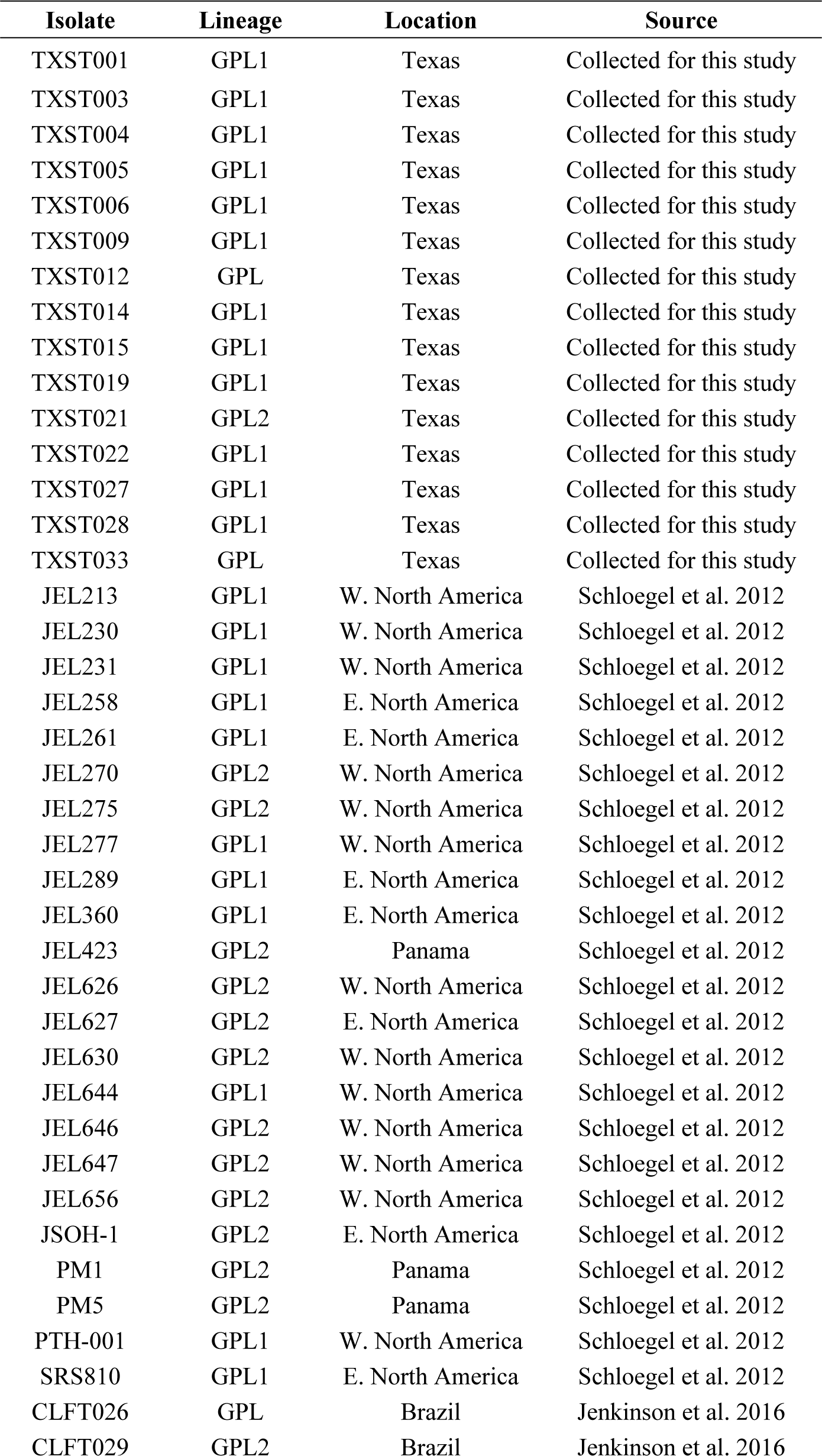
*Bd* Isolates Analyzed for this Study with Lineage, Location Collected, and Original Publication. This table includes the 68 isolates, including 15 Texas isolates, included in the STRUCTURE and pairwise *F* _ST_analyses using six MLST markers.

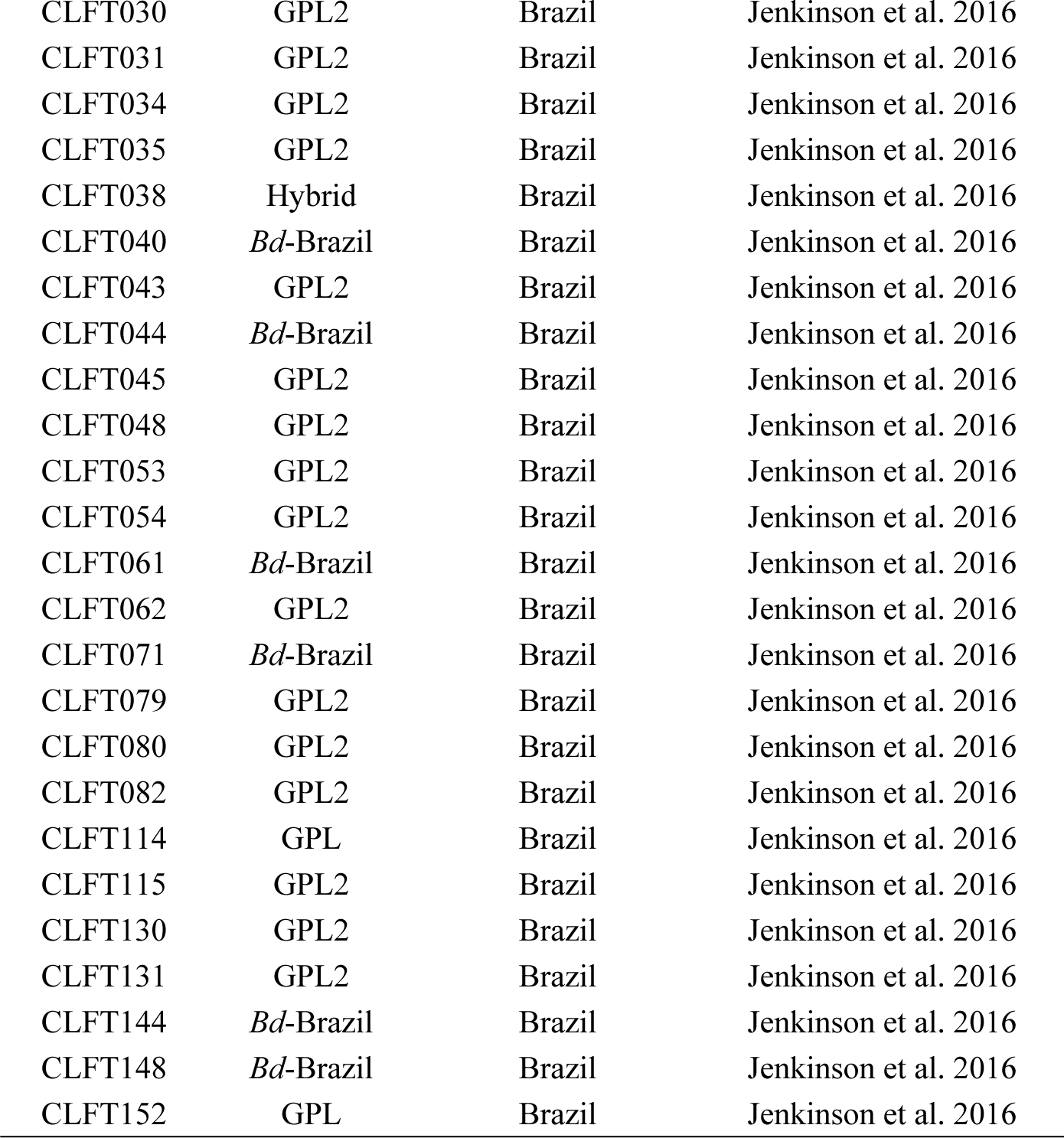

